# Impact of a caries preventive intervention in remote Indigenous Australian children

**DOI:** 10.1101/585935

**Authors:** R. Lalloo, S. K. Tadakamadla, J. Kroon, L.M. Jamieson, N. W. Johnson

## Abstract

**IMPORTANCE:** The burden of dental caries in remote Indigenous communities in Australia is unacceptably high.

**OBJECTIVES:** We tested the impact of an annual caries preventive intervention, delivered by a fly-in/fly-out professional team, on Indigenous children residing in a remote Australian community, involving selective fissure sealants, topical povidone iodine and fluoride varnish application. The outcome was caries increment at 12- and 24-month follow-up.

**DESIGN, SETTING, PARTICIPANTS:** Around 600 Indigenous children aged 5 to 17 years were invited to participate at baseline, of which 408 had caregiver consent provided. Of these, 196 consented to both the study and the treatment arm and comprised the experimental group. Two hundred and twelve consented to the epidemiological examination only, and constituted the comparison group.

**INTERVENTION:** The ‘Big Bang’ intervention, which occurred annually, comprised placement of fissure sealants, and application of povidone-iodine and fluoride varnish, following completion of each child’s dental treatment plan. Standard diet and oral hygiene advice was provided.

**MAIN OUTCOMES AND MEASURES:** Caries increment (number of tooth surfaces with new dental caries) in both primary and permanent dentitions at 12- and 24-month follow-up.

**RESULTS:** At 12-month follow-up, children in the experimental group had, on average, 5.05 (±5.47) new carious lesions compared to 7.49 (±6.94) in the comparison group (p=0.001). The preventive fraction was 33%. At 24-month follow-up, children in the experimental group had, on average, 6.47 (±6.07) new carious lesions compared to 8.43 (±5.83) in the comparison group (p=0.002). The preventive fraction was 23%.

**CONCLUSIONS AND RELEVANCE:** Indigenous children exposed to the ‘Big Bang’ caries intervention had significantly less increment in dental disease than those not exposed to the intervention. Benefits were demonstrated at both 12- and 24-month follow-ups, suggesting that the intervention is likely to be sustained if delivered across a child’s life. The cost-effectiveness of this approach is being evaluated.

*Question:* Does a quick, intensive, ‘Big Bang’ program of intervention reduce dental disease among remote Indigenous Australian children?

*Findings:* At 12- and 24-month follow-ups, children in the experimental group had fewer new caries surfaces. At the 24-month follow-up, the average caries increment was 6.47 (±6.07) in the experimental group compared to 8.43 (±5.83) in the comparison group; a preventive fraction of 23%.

*Meaning:* An intensive caries intervention delivered by a peripatetic team of professionals, involving fissure sealants, topical povidone iodine and fluoride varnish application reduces experience of dental disease among this vulnerable population.

## Introduction

Globally, oral diseases represent one of most prevalent chronic conditions and this burden has remained largely unchanged for the last 25 years.^1,2^ The number of people with untreated oral disease increased by a billion people from 2.5 in 1990 to 3.5 billion in 2015.^1^ In 2015, the total global cost due to dental conditions was more than a half-a-trillion dollars.^3^ The dominant downstream approach is making a limited impression on the burden of oral conditions at a population level. The upstream approach of addressing the fundamental social determinants and drivers of disease has enjoyed little traction.^4,5^

The National Child Oral Health Survey in Australia showed that among Aboriginal and Torres Strait Islander children (hereinafter respectfully referred to as Indigenous children) in remote communities the number of teeth in the primary dentition with decayed, missing or filled surfaces (dmfs) was 7.3, of which 5 surfaces were decayed. Almost two thirds (64%) of remote-living Indigenous children had experience of dental caries.^6^ In the permanent dentition, older remote Indigenous children had an average DMFS of 2.5, with 1.7 surfaces decayed; and 59% with caries experience.^6^ This burden is replicated in adult Australian Indigenous people;^7,8^ and across Indigenous communities globally.^9–11^

We have previously reported on a remote Indigenous community in Far North Queensland, Australia.^12^ The water supply was fluoridated across the whole community in 2005. Prior to this all school-going children were surveyed and showed that caries experience of 6- and 12-year-old children was more than twice that of the State (Queensland) average and more than four times greater than Australian children overall.^13^ A follow-up survey in this community, conducted in late 2012 by the current authors, in which over 70% of known resident schoolchildren (n=339) were examined, showed that the dental caries status had improved significantly since the pre-water fluoridation survey.12 Unfortunately the water fluoridation ceased about mid-2011.

Because caries remained a major problem, and would likely worsen without water fluoridation, our team successfully applied for funding to test the effectiveness and cost-benefit of a caries preventive intervention.^14^ This intervention was provided on an annual basis to all children who consented for the treatment phase of the study. It involved the selective placement of fissure sealants, swabbing the dentition with povidone-iodine and application of fluoride varnish (termed by us a ‘Big Bang’ intervention). At baseline, consented children received treatment for their active caries prior to the preventive intervention. Several individualised preventive interventions, such as fissure sealants, povidone-iodine and fluoride varnish, have been found to be effective in reducing caries incidence,^15–17^ but the combined effect is unknown. While fissure sealants are retained for a reasonable time and usually do not require re-application, both fluoride varnish and povidone-iodine are most effective when applied 2 to 3 times a year.^15,18^ In resource-constrained communities this is not sustainable. The aim of our study was to assess the impact of the annual ‘Big Bang’ intervention on caries increment at the 12- and 24-year follow-up visits.

## Methods

Ethics approval was granted by the Griffith University Human Research Ethics Committee (GU Ref No: DOH/05/15/HREC); the Far North Queensland Human Research Ethics Committee (FNQ HREC/15QCH/39-970); the Department of Education and Training (Queensland Government) to approach participants at the schools; and the Torres and Cape Hospital and Health Service for Site Specific Approval. Because we were interested in examining the programmatic effect of the intervention, participants were not randomly allocated to treatment and control arms. The trial was thus not registered in a clinical trial registry.

A detailed protocol of this study has been published. ^14^ All 600 school-going children (aged 5-17 years) in the community were invited to participate. At baseline in 2015, two-thirds (N=408) consented to undergo an oral epidemiological examination. All 408 were then invited to present for dental treatment (if required) through a separate consent process. The preventive intervention, comprising the placement of fissure sealants, application of povidone-iodine and fluoride varnish, followed completion of their treatment plan. Of the 408 children, 196 (experimental group) consented to the treatment and intervention phase and those who did not (n=212) formed a natural comparison group. In years 1 (2016) and 2 (2017) all children were invited to return for a clinical examination, where children in the experimental group again received the preventive intervention (Figure 1).

**Figure 1.**
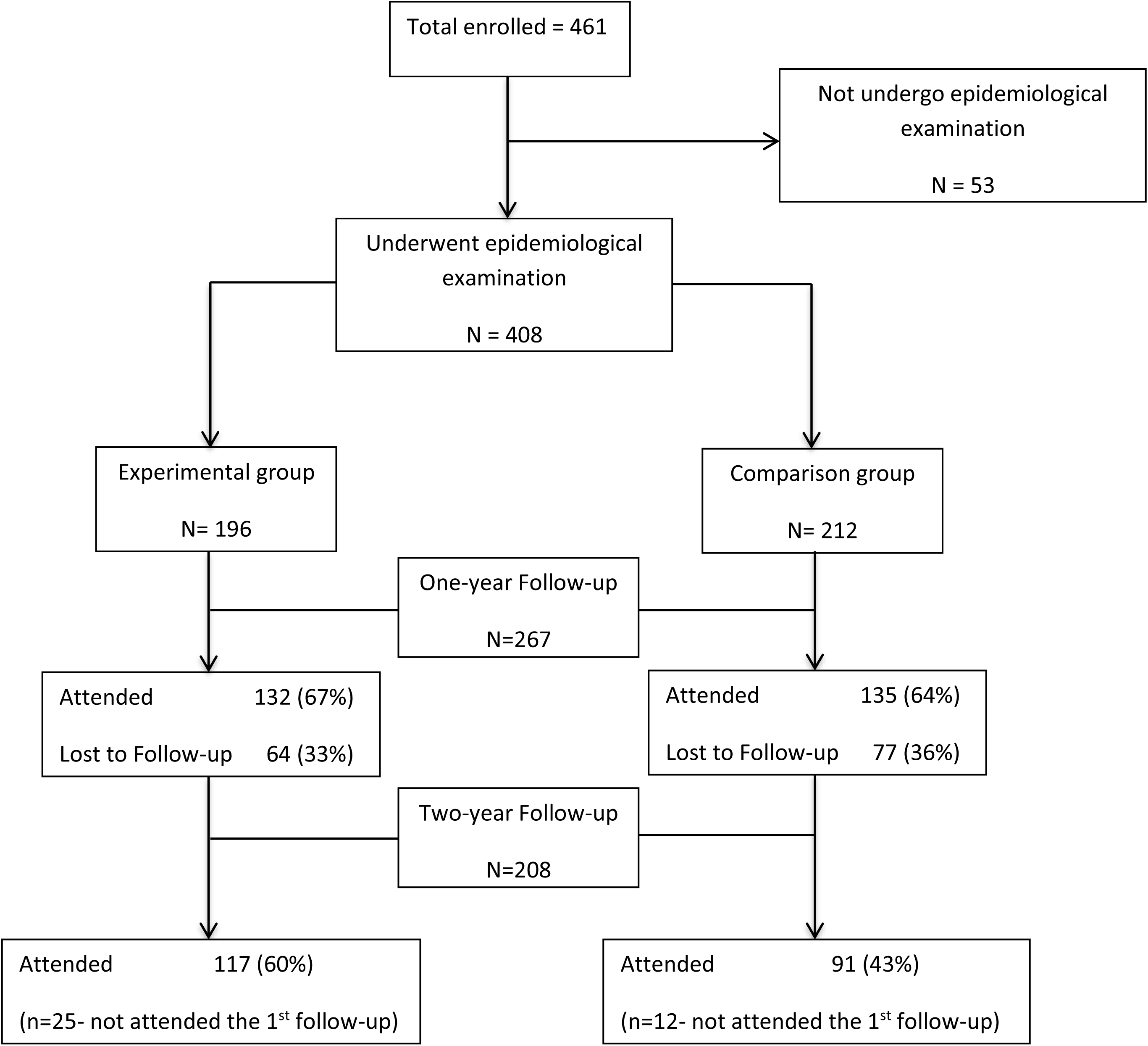
Sample size and follow-up of children by intervention and comparison groups

### Sample size calculation

A minimum sample size of 122 children (61 each in intervention and comparison group) was considered adequate to detect a clinically important difference of 30% caries reduction between the groups using a two a two-tailed z-test of proportions between two groups with 95% power and a 5% level of significance. A clinically important difference of 30% is within the 95% Confidence intervals of the prevented fraction of 0.30-0.57 and 0.24-0.51 for permanent and deciduous dentition respectively in a meta-analysis of 14 and 10 trials respectively. ^19^

The dental caries status of the children was assessed by four trained and calibrated examiners, using the International Caries Detection and Assessment system (ICDAS-II). ^20^ Children were examined in a specially set-up classroom, using a mobile, reclinable dental chair with fixed- and head-LED lights. Disposable mouth mirrors and blunt probes were used, and gauze used to control moisture. All examiners completed the ICDAS-II online training module prior to community visits. To assess inter-examiner reliability, 5% of children were re-examined by another examiner, and discrepancies in scores were discussed with the child present. The overall Kappa was 0.84, indicating a high level of agreement between examiners.

### Outcome variable

Caries increment was the outcome variable of this study. Any surface that was caries free at the baseline examination (2015) but was observed to have caries on subsequent examinations was considered as new caries, which was used for calculating the caries increment (the number of surfaces with new caries in each subject). Surfaces that were caries free at baseline examination, but were found to be extracted due to caries in subsequent examinations, were also included when calculating caries increment with all surfaces of an extracted tooth counted as missing. Caries increment was quantified for incipient (early stages of dental decay, usually identified as a white spot lesion; ICDAS 1 and 2 scores) and moderate or advanced caries (ICDAS 3 to 6) in both deciduous and permanent dentitions separately and together.

### Explanatory variables

The principal explanatory variable of this study was the group (experimental or comparison). Data were also collected on variables that could have a confounding effect, including: age and gender; daily consumption of fruit; soft drink; fruit juice; sweets & lollies; syrups and jams; and any drink with sugar added, which was recorded as yes or no. Children were asked if they brushed ‘twice or more every day’ or ‘once or less a day’. Saliva was also assessed for pH, salivary flow, buffering capacity, mutans streptococci (MS), Lactobacilli (LB) and Yeast levels using commercial chair side tests. The scoring of MS and LB loads are presented in Supplementary Figure 1. ^21^

### Statistical analysis

SPSS (IBM Corp. Released 2016. IBM SPSS Statistics for Windows, Version 24.0. Armonk, NY: IBM Corp.) was used for statistical analysis. Descriptive statistics were used to calculate mean values, standard deviations and percentages. Chi Square analysis was used to test the differences between the experimental and comparison groups for various background characteristics at baseline. As the outcome variable, caries increment, was not normally distributed, Mann-Whitney U test was used to assess the difference in the 1- and 2-year caries increment between the experimental and comparison groups. A few children in the experimental group missed the examination and intervention in 2016 (1-year follow-up) but presented at the 2-year (2017) follow-up. To assess the impact of the number of interventions (2015 and 2016 versus 2015 only) a further comparison of caries increment at the second follow-up was conducted. In order to evaluate the effect of group allocation and to adjust the impact of confounding variables, negative binomial regression analyses with log link were used. The associations between the exposure and the outcome variables are presented as Incidence Rate Ratios (IRR) with 95% confidence intervals. Four models were constructed for each caries outcome variable, the first model was unadjusted while gender and age were adjusted in the second model. In the third model, all the behavioural variables were entered along with the demographics. Finally, the fourth model comprised salivary microbial loads along with behavioural and demographic variables. A p-value of <0.05 was considered statistically significant. In order to account for attrition, an intention to treat (ITT) analysis was conducted. Caries increment at follow up were compared between the experimental and comparison groups, using both per protocol and ITT data. Unpaired t-test was used for this purpose, as the pooling methodology used for multiple imputation in SPSS does not support non-parametric tests. We calculated the preventive fraction to determine the proportion of carious lesions able to be prevented if a group was exposed to an intervention compared to a group not exposed.

Missing data were imputed using the automatic imputation method of SPSS which automatically selects fully condition specification (Markov chain Monte Carlo method) or Monotone method based on the distribution of missing data. Data were assumed to be missing at random, ten iterations were used and five imputed datasets were obtained. Linear and logistic regression were used as the univariate models for imputing scale and categorical variables respectively. Variables with complete data (group and age) were also included in the model and were used as predictors. Data were scanned to obtain the minimum and maximum values, which were then used to define constraints for each variable. Caries increment variables were rounded to the nearest integer.

## Results

At baseline, characteristics of children in the experimental arm mostly matched characteristics of their counterparts in the comparison group (Table 1). However, a higher proportion of children in the comparison group consumed soft drinks daily, added sugar to cereals and had more dental caries.

**Table 1:**
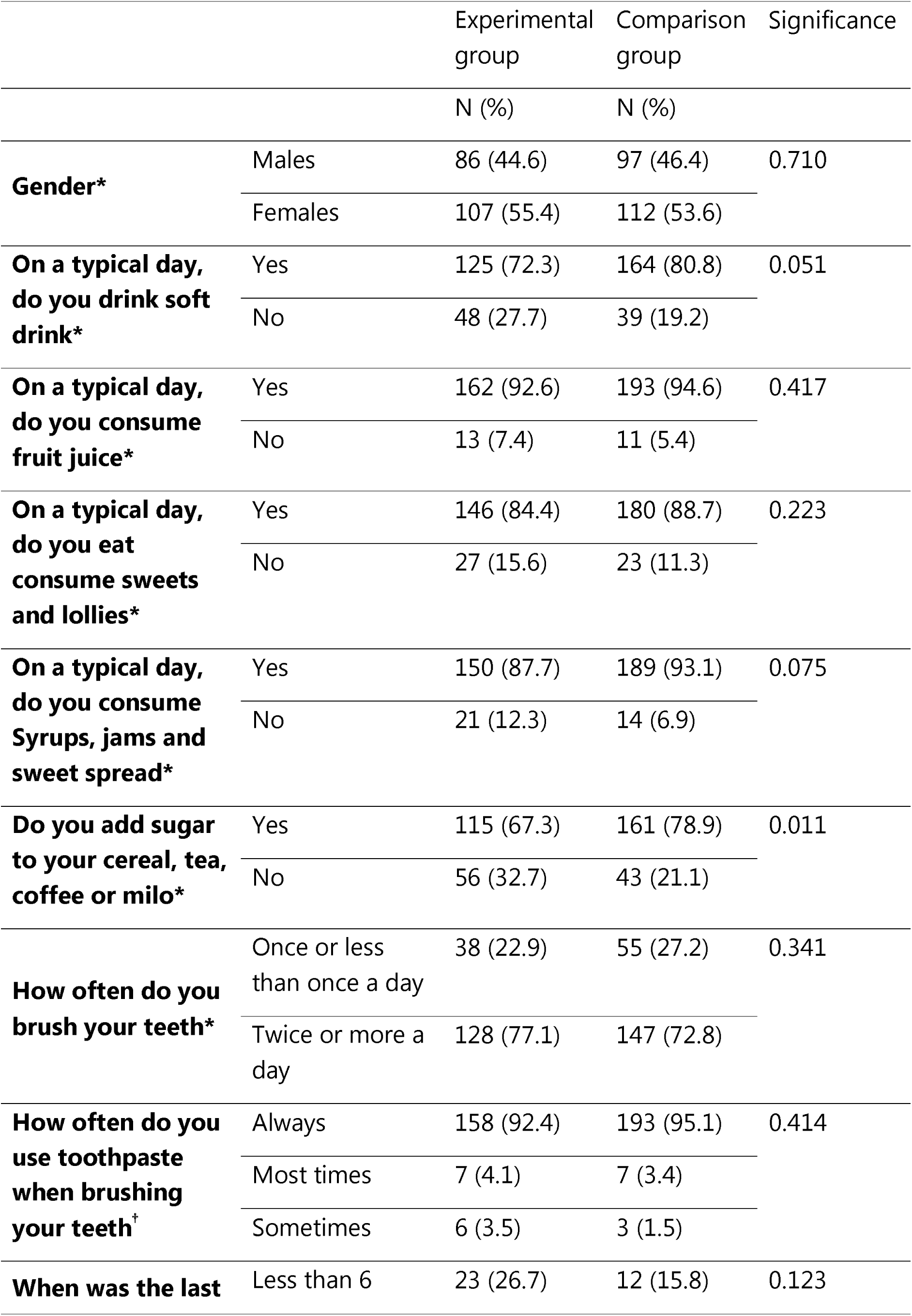
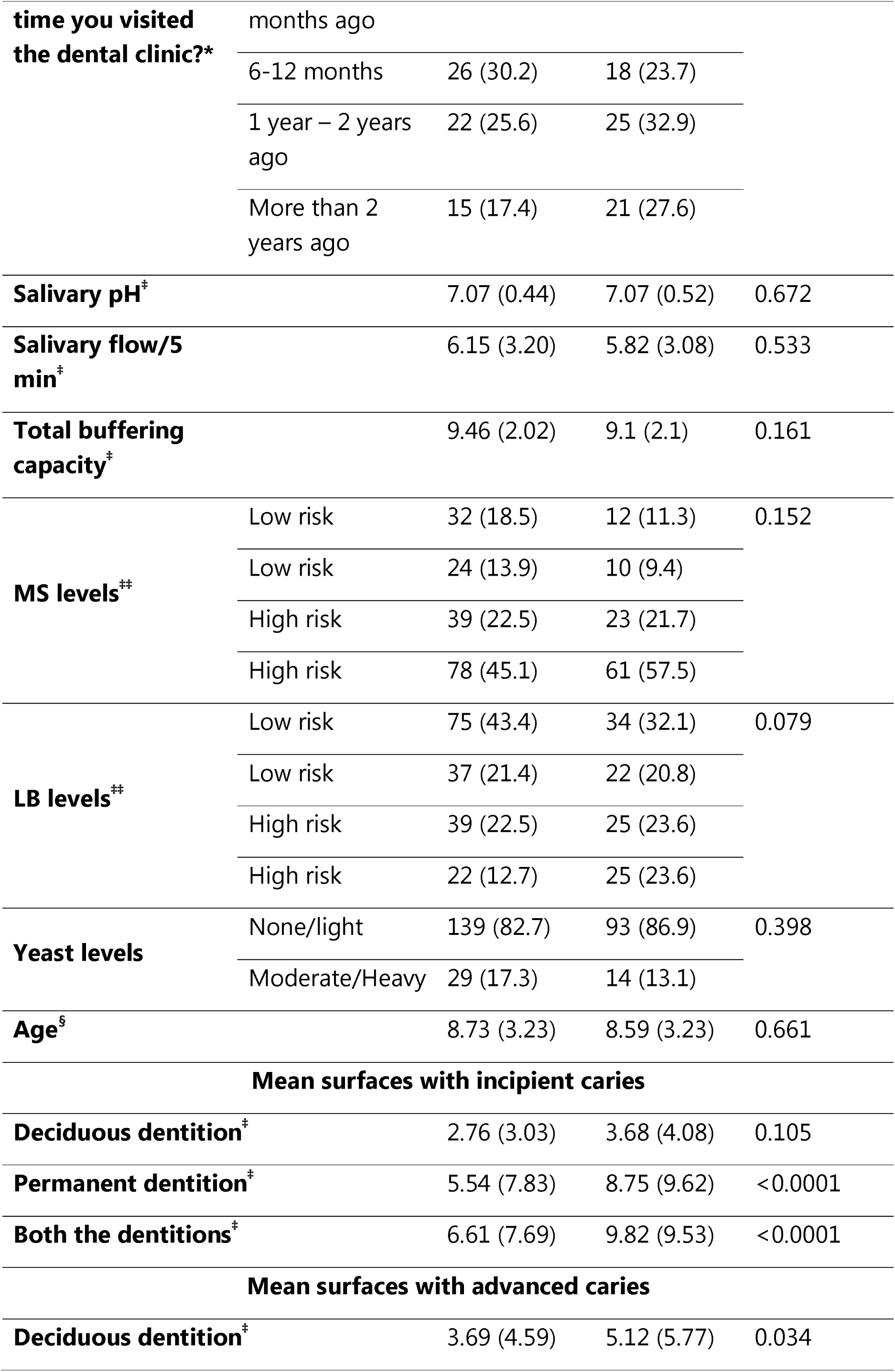
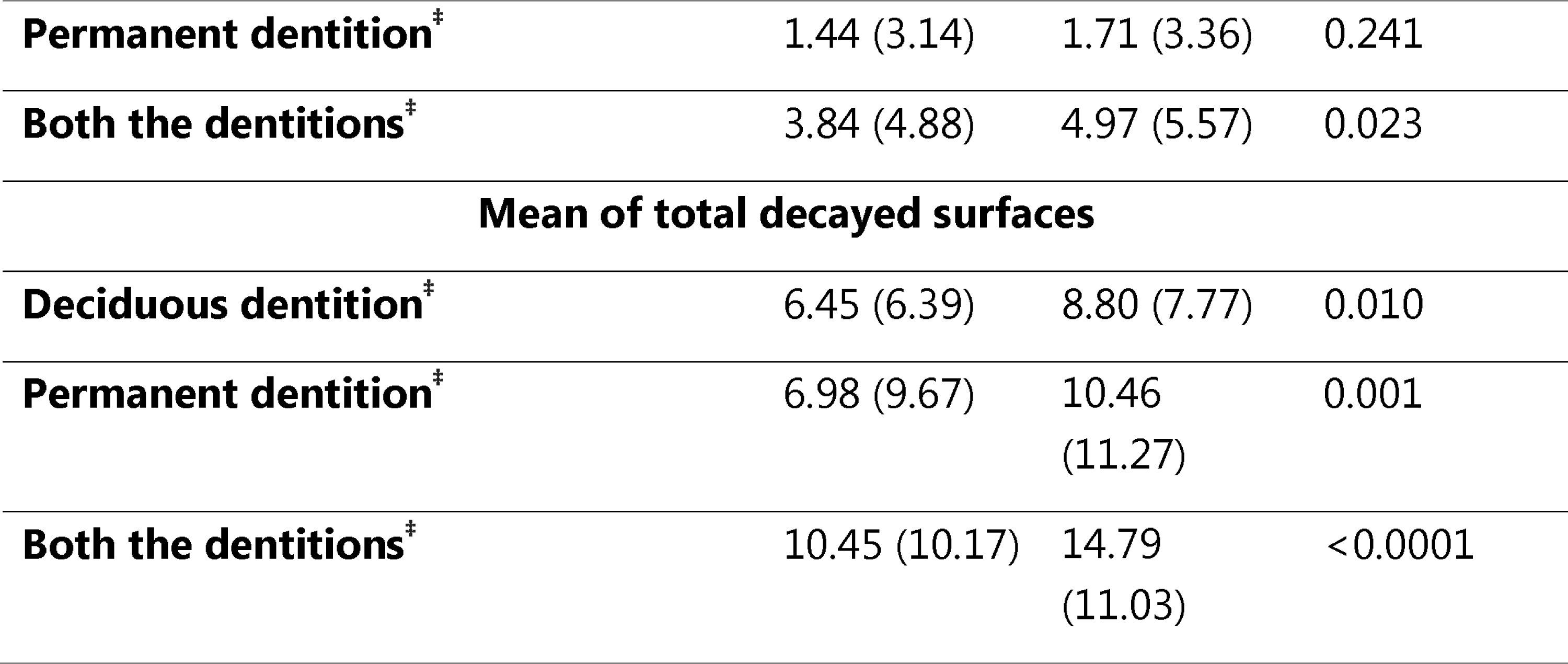
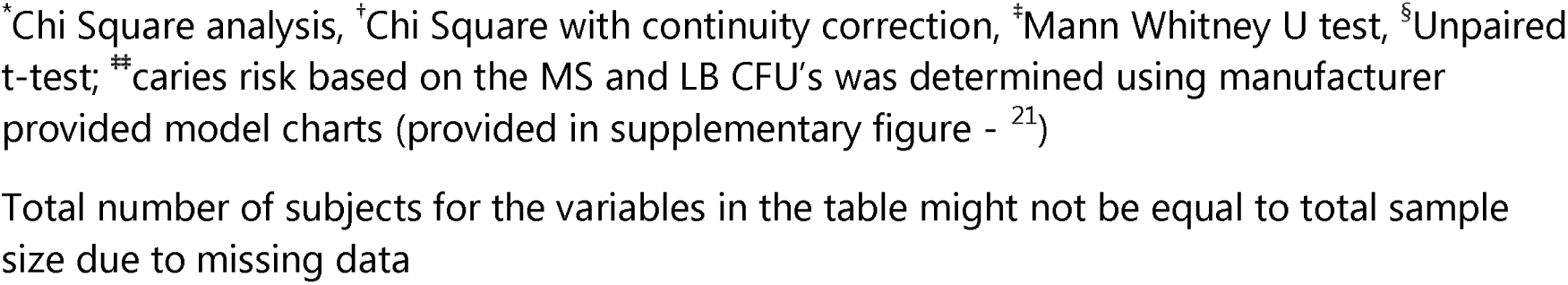
Frequency distribution of baseline characteristics between experimental and comparison groups

At the 1-year follow-up, 65% of children were retained (67% in experimental arm, 59% in comparison arm). At 2 year follow-up, 51% of children were retained (64% in experimental arm and 43% in comparison arm). (Figure 1).

At both follow-up points across all measures of caries increment, the experimental group children experienced fewer incipient and advanced caries, across both deciduous and permanent dentitions, and the differences were statistically significant in almost all comparisons (Table 2). For advanced new caries on primary tooth surfaces at the 1 year follow-up, experimental group children experienced 1.08 (±2.47) compared to 1.77 (±3.41) in the comparison group (p=0.037). In the permanent dentition, experimental group children experienced on average 3.29 (SD±3.99) new incipient carious lesions compared to 5.05 (±5.57) in the comparison group (p=0.003). Overall new caries surfaces (whole mouth) was 5.05 (±5.47) in the experimental group compared to 7.49 (±6.94) in the comparison group (p=0.001). The preventive fraction was 33% for total caries increment.

**Table 2:** Per-protocol analysis of caries increment between experimental and comparison groups

|  | N | Deciduous<br>surfaces<br>Mean (SD) | N | Permanent<br>surfaces<br>Mean (SD) | N | All<br>surfaces<br>Mean (SD) |
| --- | --- | --- | --- | --- | --- | --- |
| **1-year follow-up** |  |  |  |  |  |  |
| ***Incipient caries increment*** |  |  |  |  |  |  |
| Experimental | 90 | 1.11 (2.28) | 125 | 3.29 (3.99) | 132 | 3.87 (4.23) |
| Comparison | 83 | 1.37 (2.23) | 130 | 5.05 (5.57) | 135 | 5.71 (5.71) |
| p-value* |  | 0.130 |  | **0.003** |  | **0.004** |
| Preventive fraction<sup>†</sup> |  | 19% |  | 35% |  | 32% |
| ***Advanced caries increment*** |  |  |  |  |  |  |
| Experimental | 90 | 1.08 (2.47) | 125 | 0.46 (1.27) | 132 | 1.17 (2.49) |
| Comparison | 83 | 1.77 (3.41) | 130 | 0.71 (1.75) | 135 | 1.77 (3.27) |
| p-value* |  | **0.037** |  | 0.132 |  | **0.021** |
| Preventive fraction<sup>†</sup> |  | 39% |  | 35% |  | 34% |
| ***Total caries increment*** |  |  |  |  |  |  |
| Experimental | 90 | 2.21 (4.09) | 125 | 3.74 (4.30) | 132 | 5.05 (5.47) |
| Comparison | 83 | 3.16 (4.70) | 130 | 5.76 (6.36) | 135 | 7.49 (6.94) |
| p-value* |  | **0.036** |  | **0.005** |  | **0.001** |
| Preventive fraction<sup>†</sup> |  | 30% |  | 35% |  | 33% |
| **2-year follow-up** |  |  |  |  |  |  |
| ***Incipient caries increment*** |  |  |  |  |  |  |
| Experimental | 66 | 1.23 (1.84) | 117 | 4.37 (4.89) | 117 | 5.06 (5.30) |
| Comparison | 51 | 1.75 (2.43) | 91 | 5.37 (4.47) | 91 | 6.35 (5.01) |
| p-value* |  | 0.084 |  | **0.012** |  | **0.006** |
| Preventive fraction<sup>†</sup> |  | 30% |  | 19% |  | 20% |
| ***Advanced caries increment*** |  |  |  |  |  |  |
| Experimental | 66 | 1.53 (2.64) | 117 | 0.55 (1.31) | 117 | 1.41 (2.37) |
| Comparison | 51 | 2.00 (2.30) | 91 | 0.96 (1.44) | 91 | 2.08 (2.54) |
| p-value* |  | 0.100 |  | **0.001** |  | **0.008** |
| Preventive fraction<sup>†</sup> |  | 24% |  | 43% |  | 32% |
| ***Total caries increment*** |  |  |  |  |  |  |
| Experimental | 66 | 2.76 (3.63) | 117 | 4.91 (5.30) | 117 | 6.47 (6.07) |
| Comparison | 51 | 3.75 (3.87) | 91 | 6.33 (4.75) | 91 | 8.43 (5.83) |
| p-value* |  | 0.068 |  | **0.002** |  | **0.002** |
| Preventive fraction<sup>†</sup> |  | 26% |  | 22% |  | 23% |

\* Mann-Whitney U test, P<0.05 in bold font, <sup>†</sup>Preventive fraction = mean increment in Comparison – mean increment in Experimental ÷ mean increment in Comparison X 100

At the 2-year follow-up, children in the experimental group had, on average, 4.37 (±4.89) permanent tooth surfaces with new incipient carious lesions compared to 5.37 (±4.47) in the comparison group (p=0.012). Overall new caries surfaces in both dentitions (whole mouth) was 6.47 (±6.07) in the experimental group compared to 8.43 (±5.83) in the comparison group (p=0.002). The preventive fraction was 23% for total caries increment. In the deciduous dentition less than half of the new carious lesions were incipient. In the experimental group this was 44% compared to 47% in the comparison group. Almost all the new carious lesions in permanent teeth were incipient. In the experimental group this was 89% compared to 84% in the comparison group.

Thirty-seven participants received only one annual intervention at baseline compared with 171 children who received two interventions at baseline and the 1-year follow-up. Table 3 demonstrates that the effect at the 2-year follow-up of the intervention was more prominent in those receiving two interventions when compared to those receiving one intervention. There were no significant differences for caries increment (incipient, advanced and at both levels of caries) in the permanent dentition, and in both dentitions combined, between the experimental and comparison groups that attended only baseline and 2-year follow-up visits. However, the intervention was found to be significant, with fewer incipient lesions, advanced lesions and lesions at both levels of caries, per surface, being observed in the deciduous dentition and averaged across both dentitions, of those children who received two interventions, compared to the comparison group. When comparing only experimental group children, for all levels of carious lesions, across both dentitions, children with two interventions experienced 6.27 (±5.86) new carious surfaces compared to 7.20 (±6.87) in those who had one intervention.

**Table 3:**
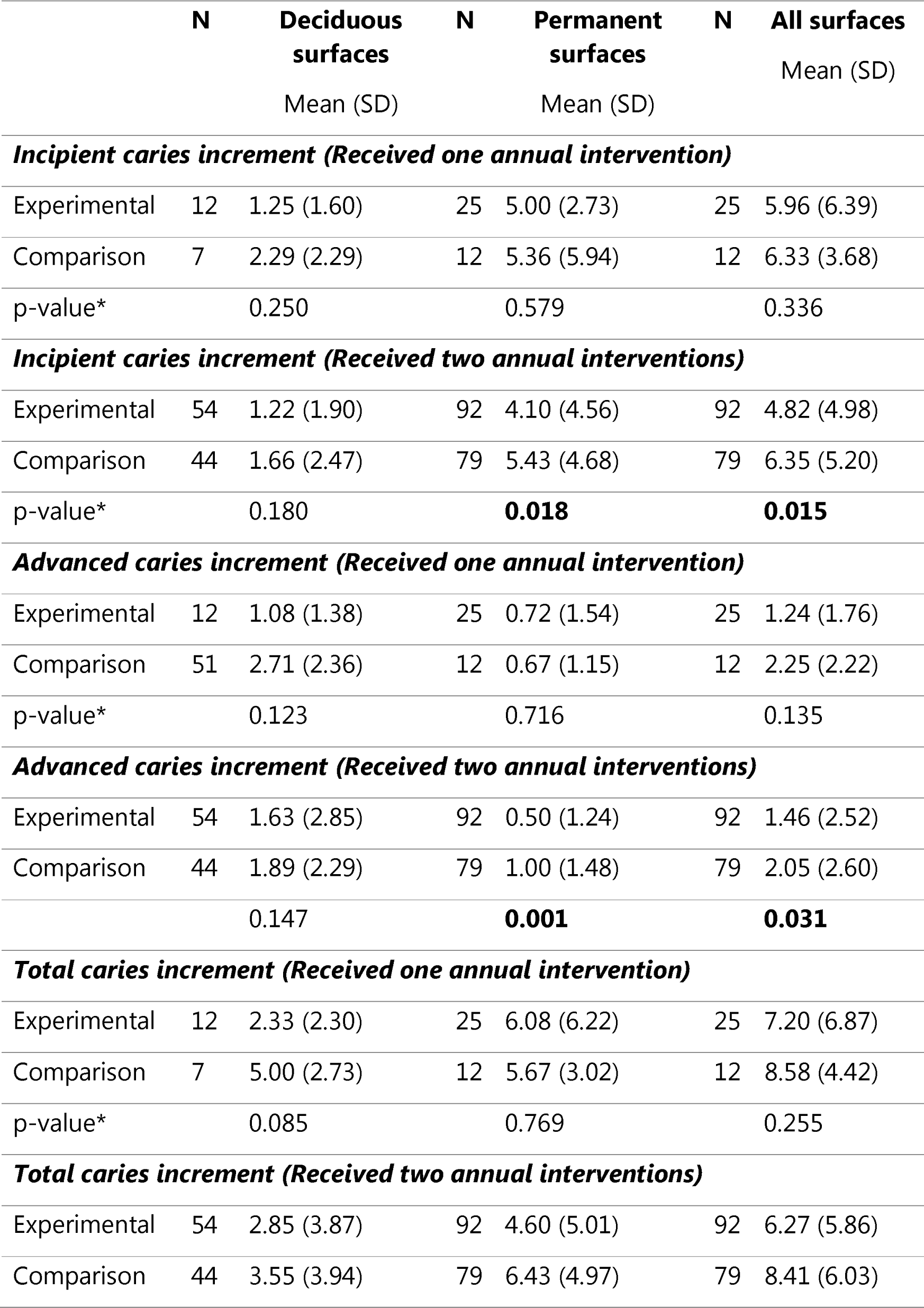
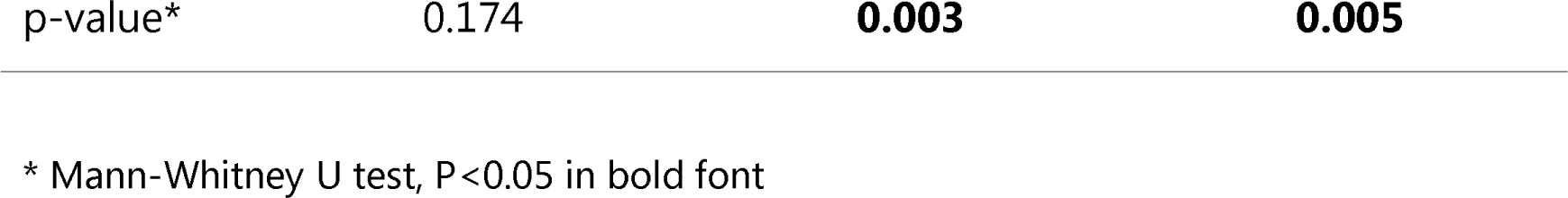
Caries increment in children attending 2-year (2017) follow-up in relation to receiving one (2015 only) and two (2015 and 2016) annual preventive interventions

|  | N | Deciduous<br>surfaces<br>Mean (SD) | N | Permanent<br>surfaces<br>Mean (SD) | N | All surfaces<br>Mean (SD) |
| --- | --- | --- | --- | --- | --- | --- |
| ***Incipient caries increment (Received one annual intervention)*** |  |  |  |  |  |  |
| Experimental | 12 | 1.25 (1.60) | 25 | 5.00 (2.73) | 25 | 5.96 (6.39) |
| Comparison | 7 | 2.29 (2.29) | 12 | 5.36 (5.94) | 12 | 6.33 (3.68) |
| p-value* |  | 0.250 |  | 0.579 |  | 0.336 |
| ***Incipient caries increment (Received two annual interventions)*** |  |  |  |  |  |  |
| Experimental | 54 | 1.22 (1.90) | 92 | 4.10 (4.56) | 92 | 4.82 (4.98) |
| Comparison | 44 | 1.66 (2.47) | 79 | 5.43 (4.68) | 79 | 6.35 (5.20) |
| p-value* |  | 0.180 |  | **0.018** |  | **0.015** |
| ***Advanced caries increment (Received one annual intervention)*** |  |  |  |  |  |  |
| Experimental | 12 | 1.08 (1.38) | 25 | 0.72 (1.54) | 25 | 1.24 (1.76) |
| Comparison | 51 | 2.71 (2.36) | 12 | 0.67 (1.15) | 12 | 2.25 (2.22) |
| p-value* |  | 0.123 |  | 0.716 |  | 0.135 |
| ***Advanced caries increment (Received two annual interventions)*** |  |  |  |  |  |  |
| Experimental | 54 | 1.63 (2.85) | 92 | 0.50 (1.24) | 92 | 1.46 (2.52) |
| Comparison | 44 | 1.89 (2.29) | 79 | 1.00 (1.48) | 79 | 2.05 (2.60) |
|  |  | 0.147 |  | **0.001** |  | **0.031** |
| ***Total caries increment (Received one annual intervention)*** |  |  |  |  |  |  |
| Experimental | 12 | 2.33 (2.30) | 25 | 6.08 (6.22) | 25 | 7.20 (6.87) |
| Comparison | 7 | 5.00 (2.73) | 12 | 5.67 (3.02) | 12 | 8.58 (4.42) |
| p-value* |  | 0.085 |  | 0.769 |  | 0.255 |
| ***Total caries increment (Received two annual interventions)*** |  |  |  |  |  |  |
| Experimental | 54 | 2.85 (3.87) | 92 | 4.60 (5.01) | 92 | 6.27 (5.86) |
| Comparison | 44 | 3.55 (3.94) | 79 | 6.43 (4.97) | 79 | 8.41 (6.03) |

|  |  |  |  |
| --- | --- | --- | --- |
| p-value* | 0.174 | **0.003** | **0.005** |

\* Mann-Whitney U test, P<0.05 in bold font

No differences were seen between the results obtained from per-protocol and ITT analyses (Table 2 and Supplemental Table 1). The preventive fraction of the intervention at the 2-year follow-up was similar in both per-protocol (23.3%) and ITT (26.2%) analyses. Caries increment was significantly different between the experimental and comparison groups in both the analyses. Therefore, the findings from the per-protocol analyses are presented in this paper. The full ITT analyses are available on request.

Table 4 and Appendix Tables 2 and 3 present the multivariable analyses of caries increment by group across the four models constructed. For the 1-year follow-up, Table 4 shows that in the full model (Model 4), comparison group children were not at a statistically increased risk of developing a new carious tooth surface compared to the experimental group children. The IRR did however range from 1.3 to 1.4 for the two dentitions combined, and for all levels of caries score. For the 2-year follow-up, children in the comparison group were at significantly greater risk (IRR=1.55; 95% CI: 1.04-2.32) to develop a new carious tooth surface in either dentition compared to experimental group children. While not statistically significant, in the two dentitions separately, the risk was substantially higher in the comparison group. The analysis by incipient (Supplemental Table 2) and advanced (Supplemental Table 3) new caries show similar findings to the overall caries experience. At the year-1 follow-up comparison group children were almost at double the risk of developing new advanced caries (IRR=1.86; 95% CI: 1.11-3.12).

**Table 4:** Multivariate analysis with total caries increment (ICDAS 1-6) as the outcome variable and group as explanatory variable

| Outcome variables | Deciduous dentition |  |  | Permanent dentition |  |  | Both the dentitions |  |  |
| --- | --- | --- | --- | --- | --- | --- | --- | --- | --- |
|  | B (SE) | IRR (95% CI) | P | B (SE) | IRR (95% CI) | P | B (SE) | IRR (95% CI) | P |
| **1- YEAR FOLLOW-UP** |  |  |  |  |  |  |  |  |  |
| **MODEL 1** |  |  |  |  |  |  |  |  |  |
| Comparison | 0.36 (0.18) | 1.43 (1.01-2.03) | **0.047** | 0.43 (0.14) | 1.54 (1.17-2.02) | **0.002** | 0.39 (0.13) | 1.48 (1.14-1.92) | **0.003** |
| Experimental | **Ref** | **Ref** |  | **Ref** | **Ref** |  | **Ref** | **Ref** |  |
| **MODEL 2** |  |  |  |  |  |  |  |  |  |
| Comparison | 0.37 (0.18) | 1.45 (1.02-2.07) | **0.038** | 0.45 (0.14) | 1.56(1.20-2.07) | **0.001** | 0.42 (0.13) | 1.51(1.17-1.97) | **0.002** |
| Experimental | **Ref** | **Ref** |  | **Ref** | **Ref** |  | **Ref** | **Ref** |  |
| **MODEL 3** |  |  |  |  |  |  |  |  |  |
| Comparison | 0.24 (0.19) | 1.27 (0.87-1.86) | 0.213 | 0.47 (0.15) | 1.60 (1.20-2.14) | **0.001** | 0.40 (0.14) | 1.50 (1.13-1.97) | **0.004** |
| Experimental | **Ref** | **Ref** |  | **Ref** | **Ref** |  | **Ref** | **Ref** |  |
| **MODEL 4** |  |  |  |  |  |  |  |  |  |
| Comparison | 0.34 (0.23) | 1.40 (0.89-2.19) | 0.144 | 0.26 (0.18) | 1.30 (0.91-1.86) | 0.149 | 0.31 (0.17) | 1.37 (0.97-1.92) | 0.072 |
| Experimental | **Ref** | **Ref** |  | **Ref** | **Ref** |  | **Ref** | **Ref** |  |
| **2- YEAR FOLLOW-UP** |  |  |  |  |  |  |  |  |  |
| **MODEL 1** |  |  |  |  |  |  |  |  |  |
| Comparison | 0.31 (0.21) | 1.36 (0.89-2.06) | 0.151 | 0.25 (0.15) | 1.29 (0.96-1.73) | 0.095 | 0.26 (0.15) | 1.30 (0.97-1.74) | 0.076 |
| Experimental | **Ref** | **Ref** |  | **Ref** | **Ref** |  | **Ref** | **Ref** |  |
| **MODEL 2** |  |  |  |  |  |  |  |  |  |
| Comparison | 0.33 (0.22) | 1.39(0.91-2.13) | 0.125 | 0.27 (0.15) | 1.31 (0.97-1.77) | 0.080 | 0.28 (0.15) | 1.32(0.98-1.78) | **0.065** |
| Experimental | **Ref** | **Ref** |  | **Ref** | **Ref** |  | **Ref** | **Ref** |  |
| **MODEL 3** |  |  |  |  |  |  |  |  |  |
| Comparison | 0.33 (0.23) | 1.40 (0.89-2.19) | 0.146 | 0.33 (0.16) | 1.39 (1.01-1.92) | **0.045** | 0.32 (0.16) | 1.37 (1.00-1.88) | **0.049** |
| Experimental | **Ref** | **Ref** |  | **Ref** | **Ref** |  | **Ref** | **Ref** |  |
| **MODEL 4** |  |  |  |  |  |  |  |  |  |
| Comparison | 0.54 (0.29) | 1.72 (0.98-3.04) | 0.061 | 0.37 (0.21) | 1.44 (0.96-2.17) | 0.080 | 0.44 (0.21) | 1.55 (1.04-2.32) | **0.033** |
| Experimental | **Ref** | **Ref** |  | **Ref** | **Ref** |  | **Ref** | **Ref** |  |

Model 1 – unadjusted; Model 2- Adjusted for age and gender; Model 3- adjusted for age, gender, behavioural and sugar intake variables (tooth brushing, fruit, soft drink, fruit juice, sweets & lollies, syrups and jams, add sugar to drink); Model 4- All the above variables +MS, LB and Yeast levels

## Discussion

The intervention to treat all existing caries at baseline followed by an annual application of selective fissure sealants, povidone-iodine and fluoride varnish was effective in reducing the number of new carious tooth surfaces in the experimental group compared to the comparison group among Indigenous children residing in a remote Australian community. At the 1- and 2-year follow-up visits the caries preventive fraction due to the intervention was in the vicinity of a third and a quarter respectively, and experimental group children experienced 2.5 and 2 fewer new carious tooth surfaces respectively.

Direct comparison of our findings to the literature is difficult as an investigation into an annual application of the interventions, to our knowledge, does not exist. The three types of intervention applied here have each been reported to have significant impact in preventing dental caries.^17,18,22–24^ Studies would usually investigate a single preventive intervention. A recent publication conducted in a rural non-fluoridated community in Chile found that bi-annual fluoride application was not effective in preschool children.^25^ A community-randomised controlled trial in Australian Aboriginal children however showed that fluoride varnish was efficacious in preventing dental caries.^26^ In this trial the caries reduction ranged from 2.3 to 3.5 surfaces per child with a preventive fraction of 24 to 36% (that is, similar to our findings).

Our study community enjoyed access to fluoridated water from 2005 to 2011, which was found to be effective in reducing the high caries burden.^12^ However this proven intervention has not been re-implemented. While the annual ‘Big Bang’ active intervention was effective, caries increment remains unacceptably high in this community, even in children who received the intervention. These interventions are resource intensive, so consideration should be given to reintroduction of water fluoridation. If active interventions are needed, consideration should be given to expanding the roles of community health workers to deliver preventive programs, especially minimally-invasive interventions such as povidone-iodine applications and fluoride varnish. In remote communities it would not be feasible or sustainable for these to be delivered by oral health professionals. If performed by trained community health workers, more regular applications as recommended rather than the annual application investigated in this study would be feasible.^27,28^

Besides the active and passive (water fluoridation) interventions, tackling social determinants is critical to reducing the burden of poor oral health. For example the consumption of sugar-sweetened beverages is of serious concern in Indigenous communities, in remote settings, with increases in consumption during adolescence.^29,30^ Strategies to reduce sugar consumption include taxes on sugar and/or sugar-sweetened beverages, and graphic warnings on labels.^31–34^

While investigating the impact of this intervention on caries increment is important, more important will be its cost utility, which is the fundamental aim of the present research program. Cost utility analyses will determine if roll-out to similar communities in Australia and globally might be recommended.

Longitudinal studies, especially in remote settings such as this one, are not without limitations. One limitation is the loss to follow-up of a significant number of participants. In our community setting there are a number of reasons for this. Firstly, school absenteeism is common and it is an important goal of government at all levels to substantially improve this.^35^ In 2015, at baseline, school attendance was high as there was a community effort to encourage children to attend school. In the follow-up visits this effort had waned and absenteeism rates were higher. Some children, especially later into their schooling, move to larger towns and cities to complete their education. Significant community events affect school attendance. While we lost a number of children over the 2 years of the study, the per-protocol and ITT analysis showed no significant differences. We believe this loss to follow-up, while regrettable, does not negate the findings of the study. Responses for (as an example) number of times children brushed may be over-estimated and was not verified by any parental information. While clinical examiners were not informed on the group status of the children who attended follow-up examinations, the presence of fissures sealants made blinding impossible.

While it is important to address the social determinants of health in Indigenous communities in Australia, it is critically important that at a national level there is progress and resolution on the broader issues of Indigenous disadvantage and dispossession of land and resources. Many argue for formal recognition of the Aboriginal and Torres Strait Islander peoples in the Nation’s constitution, for a formal treaty to acknowledge and address the impact of colonisation and a Makarrata: *the coming together after a struggle*,^36,37^ all of which impact on oral health, oral health-related quality of life and overall health and well-being.

## Conclusion

Remote Indigenous children who received the annual ‘Big Bang’ preventive intervention experienced significantly fewer carious lesions at 12- and 24-month follow-ups, compared to children who did not receive the intervention. However, high levels of dental disease were still experienced by both groups. It is incumbent upon all dental researchers, dental providers and policy makers to advocate for public health interventions of known cost-effectiveness to reduce these unacceptable oral health inequalities.

## Author Contributions

R. Lalloo, contributed to design, data acquisition, analysis, and interpretation, drafted and critically revised the manuscript; S. Kumar, contributed to data analysis and interpretation, and critically revised the manuscript; N.W. Johnson, and J Kroon contributed to design, data acquisition, data interpretation and critically revised the manuscript. L Jamieson critically revised the manuscript. All authors gave final approval and agree to be accountable for all aspects of the work.

## Acknowledgements

The authors gratefully acknowledge the Elders, Community Members & Community Workers in the Northern Peninsula Area of Far North Queensland and the Principals, Staff & Children of the Northern Peninsula Area State College. Our sincerest thank you to all Chief and Associate Investigators and Project Managers. This study was funded by the Australian National Health and Medical Research Council (NHMRC, Project Grant APP1081320). The authors declare no potential conflicts of interest. NWJ had full access to all the data in the study and takes responsibility for the integrity of the data and the accuracy of the data analysis.

## Access to Data

Data is available upon request to the corresponding author.

---

**Supplementary Figure 1.**
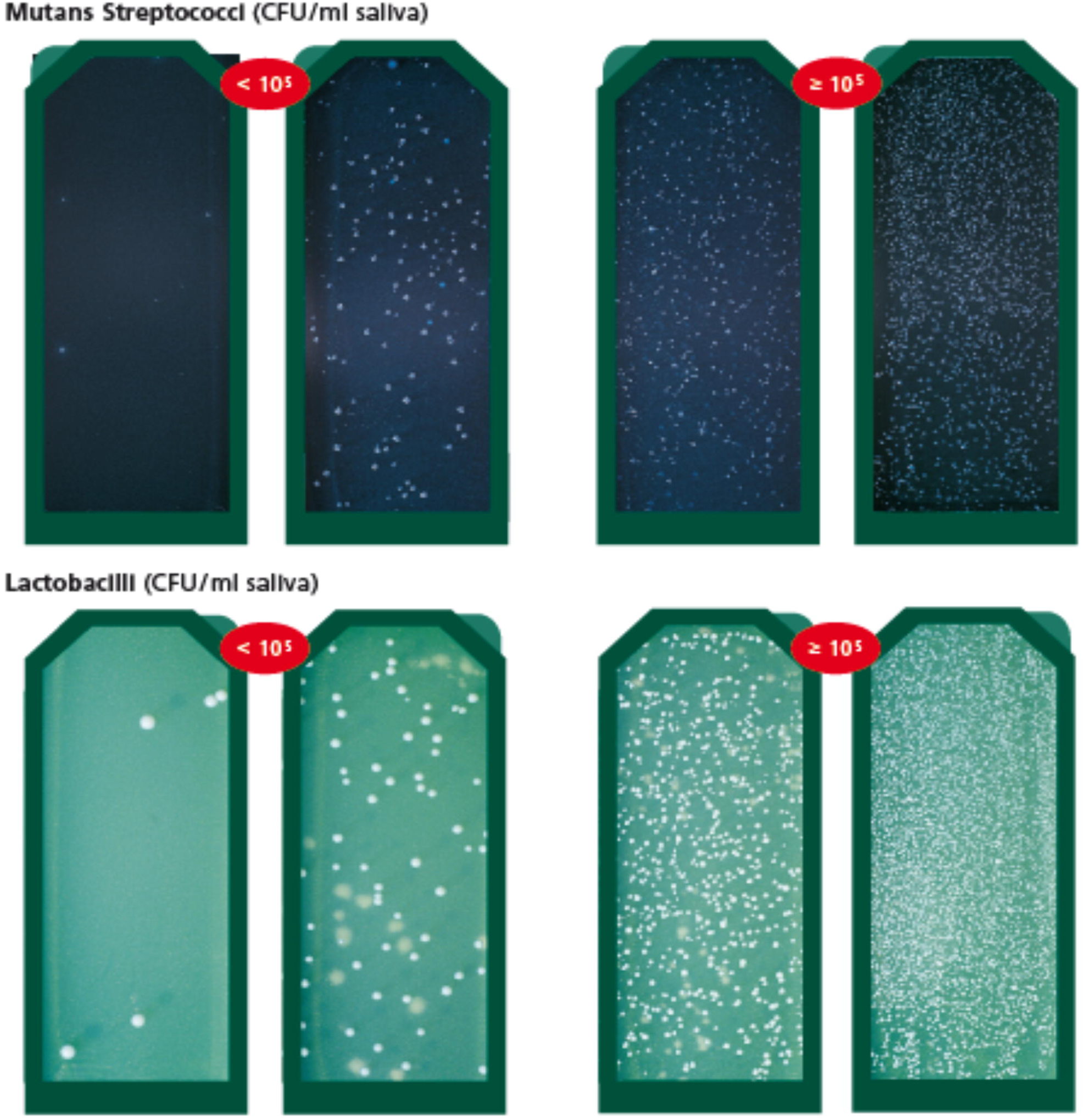
Scoring of MS and LB counts (CFU/ml saliva)

