## Supplementary material for "Impact of a caries preventive intervention in remote Indigenous Australian children": Supplemenatry tables

***Supplemental Table 1: Intention to Treat analysis of caries increment between experimental and comparison groups***

|  | | N | Deciduous surfaces  Mean (SE) | Permanent surfaces  Mean (SE) | All surfaces  Mean (SE) |
| --- | --- | --- | --- | --- | --- |
| *1-year follow-up* | | | | | |
| *Incipient caries increment* | | | | | |
| Experimental | | 196 | 1.73 (0.21) | 4.17 (0.38) | 5.90 (0.44) |
| Comparison | | 212 | 2.27 (0.37) | 5.51 (0.40) | 7.79 (0.47) |
| p-value* | |  | 0.166 | **0.018** | **0.005** |
| Preventive fraction^†^ | |  | 24% | 24% | 24% |
| *Advanced caries increment* | | | | | |
| Experimental | | 196 | 2.08 (0.35) | 0.82 (0.13) | 2.90 (0.37) |
| Comparison | | 212 | 2.77 (0.27) | 1.02 (0.15) | 3.79 (0.30) |
| p-value* | |  | **0.058** | 0.132 | **0.027** |
| Preventive fraction^†^ | |  | 25% | 20% | 24% |
| *Total caries increment* | | | | | |
| Experimental | | 196 | 3.81 (0.41) | 4.99 (0.45) | 8.80 (0.54) |
| Comparison | | 212 | 5.05 (0.50) | 6.53 (0.50) | 11.58 (0.60) |
| p-value* |  |  | **0.009** | **0.018** | **<0.0001** |
| Preventive fraction^†^ |  |  | 25% | 24% | 24% |
| *2-year follow-up* | | | | | |
| *Incipient caries increment* | | | | | |
| Experimental | | 196 | 2.28 (0.23) | 5.17 (0.37) | 7.45 (0.50) |
| Comparison | | 212 | 2.83 (0.48) | 6.10 (0.51) | 8.93 (0.69) |
| p-value* |  |  | 0.244 | 0.191 | 0.096 |
| Preventive fraction^†^ |  |  | 19% | 15% | 17% |
| *Advanced caries increment* | | | | | |
| Experimental | | 196 | 2.44 (0.28) | 0.82 (0.12) | 3.26 (0.27) |
| Comparison | | 212 | 2.80 (0.66) | 1.40 (0.11) | 4.19 (0.68) |
| p-value* |  |  | 0.567 | **0.003** | 0.175 |
| Preventive fraction^†^ |  |  | 13% | 41% | 22% |
| *Total caries increment* | | | | | |
| Experimental | | 196 | 4.72 (0.45) | 5.99 (0.43) | 10.71 (0.65) |
| Comparison | | 212 | 5.63 (1.06) | 7.50 (0.51) | 13.13 (1.08) |
| p-value* |  |  | 0.364 | **0.053** | **0.042** |
| Preventive fraction^†^ |  |  | 16% | 20% | 18% |

* Unpaired t-test, P<0.05 in bold font, ^†^Preventive fraction = mean increment in Comparison – mean increment in Experimental ÷ mean increment in Comparison X 100

***Supplemental Table 2: Multivariate analysis with incipient caries increment (ICDAS 1-2) as the outcome variable and group as explanatory variable***

|  | **Model 1** | | | **Model 2** | | | **Model 3** | | | **Model 4** | | |
| --- | --- | --- | --- | --- | --- | --- | --- | --- | --- | --- | --- | --- |
| **Outcome variables** | B (SE) | IRR (95% CI) | P | B (SE) | IRR (95% CI) | P | B (SE) | IRR (95% CI) | P | B (SE) | IRR (95% CI) | P |
| ***1-year follow-up*** | | | | | | | | | | | | |
| **Incipient caries increment in the Deciduous dentition** | | | | | | | | | |  |  |  |
| Comparison | 0.21 (0.20) | 1.24 (0.83-1.85) | 0.301 | 0.24 (0.21) | 1.27 (0.85-1.90) | 0.254 | 0.07 (0.22) | 1.07 (0.70-1.65) | 0.750 | 0.07 (0.26) | 1.08(0.65-1.79) | 0.777 |
| Experimental | **Ref** | **Ref** |  | **Ref** | **Ref** |  | **Ref** | **Ref** |  | **Ref** | **Ref** |  |
| **Incipient caries increment in the permanent dentition** | | | | | | | | | |  |  |  |
| Comparison | 0.43 (0.14) | 1.54 (1.17-2.02) | **0.002** | 0.47 (0.14) | 1.60(1.21-2.11) | **0.001** | 0.50 (0.15) | 1.65 (1.23-2.21) | **0.001** | 0.27 (0.18) | 1.31 (0.91-1.88) | 0.141 |
| Experimental | **Ref** | **Ref** |  | **Ref** | **Ref** |  | **Ref** | **Ref** |  | **Ref** | **Ref** |  |
| **Incipient caries increment in both the dentitions** | | | | | | | | | |  |  |  |
| Comparison | 0.39 (0.14) | 1.48 (1.13-1.92) | **0.004** | 0.42 (0.14) | 1.53(1.17-1.99) | **0.002** | 0.43 (0.14) | 1.54 (1.16-2.05) | **0.003** | 0.26 (0.18) | 1.29 (0.91-1.83) | 0.147 |
| Experimental | **Ref** | **Ref** |  | **Ref** | **Ref** |  | **Ref** | **Ref** |  | **Ref** | **Ref** |  |
| ***2-year follow-up*** | | | | | | | | | | | | |
| **Incipient caries increment in the Deciduous dentition** | | | | | | | | | |  |  |  |
| Comparison | 0.35 (0.24) | 1.42 (0.89-2.28) | 0.145 | 0.38 (0.24) | 1.46(0.90-2.35) | 0.122 | 0.32 (0.26) | 1.34 (0.83-2.28) | 0.217 | 0.61 (0.32) | 1.84 (0.98-3.44) | **0.057** |
| Experimental | **Ref** | **Ref** |  | **Ref** | **Ref** |  | **Ref** | **Ref** |  | **Ref** | **Ref** |  |
| **Incipient caries increment in the permanent dentition** | | | | | | | | | |  |  |  |
| Comparison | 0.21 (0.15) | 1.23 (0.91-1.66) | 0.177 | 0.23 (0.16) | 1.26(0.93-1.71) | 0.140 | 0.29 (0.17) | 1.34 (0.97-1.86) | **0.077** | 0.35 (0.21) | 1.41 (0.94-2.14) | 0.101 |
| Experimental | **Ref** | **Ref** |  | **Ref** | **Ref** |  | **Ref** | **Ref** |  | **Ref** | **Ref** |  |
| **Incipient caries increment in both the dentitions** | | | | | | | | | |  |  |  |
| Comparison | 0.23 (0.15) | 1.26 (0.93-1.69) | 0.133 | 0.25 (0.15) | 1.28(0.95-1.73) | 0.106 | 0.29 (0.16) | 1.34 (0.97-1.84) | 0.078 | 0.41 (0.21) | 1.50 (1.00-2.26) | **0.05** |
| Experimental | **Ref** | **Ref** |  | **Ref** | **Ref** |  | **Ref** | **Ref** |  | **Ref** | **Ref** |  |

Model 1 – unadjusted; Model 2- Adjusted for age and gender; Model 3- adjusted for age, gender, behavioural and sugar intake variables (tooth brushing, fruit, soft drink, fruit juice, sweets & lollies, syrups and jams, add sugar to drink); Model 4- All the above variables +MS, LB and Yeast levels

***Supplemental Table 3: Multivariate analysis with advanced caries increment (ICDAS 3-6) as the outcome variable and group as explanatory variable***

|  | **Model 1** | | | **Model 2** | | | **Model 3** | | | **Model 4** | | |
| --- | --- | --- | --- | --- | --- | --- | --- | --- | --- | --- | --- | --- |
| **Outcome variables** | B (SE) | IRR (95% CI) | P | B (SE) | IRR (95% CI) | P | B (SE) | IRR (95% CI) | P | B (SE) | IRR (95% CI) | P |
| ***1-year follow-up*** | | | | | | | | | | | | |
| **Advanced caries increment in the deciduous dentition** | | | | | | | | | |  |  |  |
| Comparison | 0.50 (0.20) | 1.64 (1.11-2.44) | **0.013** | 0.51 (0.20) | 1.67 (1.12-2.47) | **0.011** | 0.43 (0.22) | 1.53 (0.99-2.37) | **0.053** | 0.62 (0.26) | 1.86(1.11-3.12) | **0.019** |
| Experimental | **Ref** | **Ref** |  | **Ref** | **Ref** |  | **Ref** | **Ref** |  | **Ref** | **Ref** |  |
| **Advanced caries increment in the permanent dentition** | | | | | | | | | |  |  |  |
| Comparison | 0.44 (0.21) | 1.55 (1.03-2.34) | **0.036** | 0.35 (0.21) | 1.42(0.94-2.16) | 0.099 | 0.28 (0.22) | 1.32 (0.86-2.03) | **0.205** | 0.20 (0.27) | 1.23 (0.73-2.07) | 0.445 |
| Experimental | **Ref** | **Ref** |  | **Ref** | **Ref** |  | **Ref** | **Ref** |  | **Ref** | **Ref** |  |
| **Advanced caries increment in both the dentitions** | | | | | | | | | |  |  |  |
| Comparison | 0.42 (0.16) | 1.52 (1.11-2.08) | **0.009** | 0.39 (0.16) | 1.48(1.08-2.03) | **0.015** | 0.30 (0.17) | 1.36 (0.97-1.90) | **0.079** | 0.50 (0.21) | 1.64 (1.09-2.48) | **0.018** |
| Experimental | **Ref** | **Ref** |  | **Ref** | **Ref** |  | **Ref** | **Ref** |  | **Ref** | **Ref** |  |
| ***2-year follow-up*** | | | | | | | | | | | | |
| **Advanced caries increment in the deciduous dentition** | | | | | | | | | |  |  |  |
| Comparison | 0.27 (0.23) | 1.31 (0.83-2.07) | 0.251 | 0.29 (0.24) | 1.34(0.84-2.13) | 0.215 | 0.35 (0.25) | 1.41 (0.86-2.31) | 0.169 | 0.47 (0.32) | 1.61 (0.85-3.03) | 0.143 |
| Experimental | **Ref** | **Ref** |  | **Ref** | **Ref** |  | **Ref** | **Ref** |  | **Ref** | **Ref** |  |
| **Advanced caries increment in the permanent dentition** | | | | | | | | | |  |  |  |
| Comparison | 0.56 (0.22) | 1.75 (1.15-2.67) | **0.010** | 0.53 (0.22) | 1.70 (1.11-2.61) | **0.015** | 0.55 (0.23) | 1.73 (1.10-2.71) | **0.018** | 0.48 (0.29) | 1.62 (0.93-2.85) | 0.092 |
| Experimental | **Ref** | **Ref** |  | **Ref** | **Ref** |  | **Ref** | **Ref** |  | **Ref** | **Ref** |  |
| **Advanced caries increment in both the dentitions** | | | | | | | | | |  |  |  |
| Comparison | 0.39 (0.18) | 1.47 (1.04-2.08) | **0.028** | 0.38 (0.18) | 1.46(1.03-2.07) | **0.033** | 0.40 (0.19) | 1.49 (1.03-2.16) | **0.034** | 0.54 (0.24) | 1.72 (1.07-2.77) | **0.025** |
| Experimental | **Ref** | **Ref** |  | **Ref** | **Ref** |  | **Ref** | **Ref** |  | **Ref** | **Ref** |  |

Model 1 – unadjusted; Model 2- Adjusted for age and gender; Model 3- adjusted for age, gender, behavioural and sugar intake variables (tooth brushing, fruit, soft drink, fruit juice, sweets & lollies, syrups and jams, add sugar to drink); Model 4- All the above variables +MS, LB and Yeast levels
